# Different profile of transcriptome between wheat Yunong 201 and its high-yield mutant Yunong 3114

**DOI:** 10.1101/005496

**Authors:** Feng Chen, Zhongdong Dong, Zhang Ning, Zhang Xiangfen, Dangqun Cui

## Abstract

Wheat is one of the most important crops in the world. With the exponentially increasing population and the need for ever increased food and feed production, an increased yield of wheat grain (as well as rice, maize and other grains) will be critical. Modern technologies are utilized to assist breeding programs. Such as the transcriptome sequencing, which greatly improves our genetic understanding, provides a platform for functional genomics research on crops. Herein, to get an overview of transcriptome characteristics of Yunong 3114, which is screened from the EMS mutagenized population of, a high quality Chinese winter noodle wheat, due to its different plant architecture as well as larger kernel size and higher grain weight, a high-throughput RNA sequencing based on next generation sequencing technology (Illumina) were performed. These unigenes were annotated by Blastx alignment against the NCBI non-redundant (nr), Clusters of orthologous groups (COG), gene orthology (GO), and the Kyoto Encyclopedia of Genesand Genomes (KEGG) databases. The 90.96% of the unigenes matched with protein in the NCBI nr database. Functional analysis identified that changes in several GO categories, including recognition of pollen, apoptotic process, defense response, receptor activity, protein kinase activity, DNA integration and so forth, played crucial roles in the high-yield characteristics of the mutant. Real-time PCR analysis revealed that the recognition of pollen related gene GsSRK is significantly up-regulated in Yunong 3114. In addition, alternative splicing (AS) analysis results indicated that mutation influence AS ratio, especially the retained introns, including the pollen related genes. Furthermore, the digital gene expression spectrum (DGE) profiling data provides comprehensive information at the transcriptional level that facilitates our understanding of the molecular mechanisms of various physiological aspects including development and high-yield of wheat. Together, these studies substantially increase our knowledge of potential genes and pathways for the genetic improvement of wheat and provide new insights into the yield and breeding strategies.

## Introduction

Wheat (*Triticum aestivum* L.) is one of the world’s most important food staple sources, which nowadays is grown a broad range of climates throughout the world. To the best of our knowledge, wheat was domesticated to provide calories and protein for human-kind from approximately 10,000 years ago and has been adapted and ranked with rice and maize as the world’s most cultivated crop plants [1]. In the year of 2011, the total world production of wheat reached approximately 702×10^6^ metric tons, with rice ∼ 723×10^6^ tons and maize ∼ 885×10^6^ tons [2]. To support the several billion people living on the earth, the production of high-quality food must increase with reduced inputs. Many modern biotechnologies are exploited to assist wheat breeding programs to increase crop yield, as well as nutritional content, drought and salinity tolerance and biotic tolerance [3–6]. Grain number, panicle number, and grain weight are three important components of grain yield of wheat. Grain weight is largely influenced by grain size, which is determined by its three dimensions (length, width, and thickness) and the degree of filling [7]. Improvement of these factors plays a pivotal role in further yield increase in wheat-breeding programs.

Molecular understanding of the biology of the developing grain and the knowledge of the full genome sequence of wheat will undeniably highly assist the wheat improvement of yield and quality traits [4, 8]. High quality transcriptomics analysis method increased this understanding. For example, wheat transcriptome sequencing can be performed to identify candidate genes for a certain phenotype and to develop SNP markers for tracking favourable alleles in breeding programs [9, 10]. Knowledge of the transcriptome also provides great aids for the design of microarrays and the interpretation of RNA-Seq experiments [11, 12].

Assembling the sequence of the transcriptome, in conjunction with the genome sequence analysis, will promote us to gain insight into the plant’s capacity for high-level transient gene expression, generation of mobile gene silencing signals, and hyper-susceptibility to viral infection and environment [13–16]. The difficulty of sequence assembly is the most significant factor explaining the relatively small number of plant species with finished genome sequences. Plant genomes tend to be large and highly repetitive; they have a propensity to contain large gene families and are frequently polyploid. For example, wheat is allohexaploid with a large genome, estimated at 17 Gb roughly 5 times the size of the human genome [17, 18]. Additionally, the wheat genome is estimated to be 90% repetitive [19], which is a significantly higher percentage than Brachypodium (22%), rice (26%), and sorghum (54%) [20]. Thus, the large size and the complex polyploid nature of the wheat genome make genetic and functional analysis extremely challenging.

To assemble such large and complex genomes, exceedingly efficient computational procedure with both time and memory resources is essential, simultaneously highly accurate to avoid mis-assembly of closely related sequences is also a requirement [4]. Recently, *de novo* transcript assembly analysis has been used to comprehensively analyze transcriptomes. Several recent studies were performed using massively parallel sequencing technology, to detail the transcriptome sequencing of various non-model species, including wheat [21, 22]. *De novo* assembly of short sequences of transcripts enables researchers to reconstruct the sequences of entire transcriptomes, identify and catalogue all of the expressed genes, separate isoforms, and capture transcript expression levels [23].

The Chinese winter wheat cultivar Yunong 201, showing outstanding dry white noodle quality and released in 2006 as a high-quality noodle wheat cultivar. In this study, we got a high yield cultivar line Yunong 3114 derived from Yunong 201, with a significant increase in grain length, thousand seed weight and plot yield. To identify putative effector genes, transcriptome sequencing (RNA-Seq) is now being adopted to analyze the transcriptomes of Yunong 201 and Yunong 3114, in conjunction with Trinity for assembling. Based on our transcriptome data, a digital gene expression (DGE) system was used to compare the gene expression profiles of Yunong 3114 and 201. The unigenes were then annotated to putative functions, classifications or pathways by sequence similarity analysis against the sequences in the public database resources. Of these, 89,644 unigenes abundance were observed which were categorised into 3 terms in GO database, listing as biological process, cellular components and molecular function. Using a 2-fold cutoff, we identified a large changes in enriched expression unigenes relating to primary metabolic pathways, DNA synthesis, ATP binding, recognition of pollen, apoptotic process, defense response and so forth. Alternative splicing (AS) play important roles in increasing protein variation and complexity, function diversities and regulating gene expression [24]. We also analyzed the AS condition in Yunong 3114 and Yunong 201 to confirm if mutation may change AS ratio. The results will allow us to reveal the molecular mechanism underlying the different yield phenotypes of different genotypes, and may facilitate better understanding the pathways and genes associated with yield improvement of wheat.

## Materials and Methods

### Plant growth and treatment

A Chinese winter wheat cultivar Yunong 201, showing outstanding dry white noodle quality and released in 2006 as a high-quality noodle wheat cultivar, was treated by EMS (ethyl methanesulfonate). A M_2_ line was screened from the 0.8% EMS mutagenized population of Yunong 201 due to its different plant architecture as well as larger kernel size and higher grain weight, and was self-crossed for four times into a M_6_ line Yunong 3114. The high-yield Yunong 3114 showed a significant increase in grain length (7.5 cm / 10 kernels) and thousand seed weight (45.1 g), whereas 10-grain length and thousand seed weight of wild-type Yunong 201 are 6.4 cm and 39.7 g, respectively. Yunong 201 and its derived line Yunong 3114 were planted and grown at the Zhengzhou Scientific Research and Education Center of Henan Agricultural University during 2011-12 cropping seasons under non-stressed conditions.

The flowers and spikes samples were collected from both lines at 7 days, 14 days, 21 days, 28 days and 35 days after flowering, and mature seeds (deposit 1 year), respectively, and used for generation of small RNA libraries of Yunong 201 and Yunong 3114.

## Transcriptome Sequencing and Quality filtering

The Solexa RNA paired-end sequencing was performed to generate a number of reads for transcriptome sequencing, with sequencing performed by commercial service provider (Oebiotech, Shanghai, China).

Quality trimming of raw Solexa reads was performed using custom Perl scripts according to the following prescription: a) the sliding window method to remove low quality: the quality threshold 20 (e 1% -O 5 bp, length was < 35 bp; b) removal in sequence containing N parts: reads were discarded if the length was <35 bp. The assembly was done using standard settings.

### *De novo* assembly

To generate a non-redundant set of transcripts, we performed a *de novo* assembly with effective wheat solexa sequencing reads, using the software Trinity (ver. trinityrnaseq_r2012-10-05, paired-end assembly method). Trinity, a short reads assembling program developed specifically for *de novo* transcriptome assembly from short-read RNA-Seq data.

The relatively long-read sequence data were obtained with the 454 sequencing system. In addition, contig construction is greatly affected by sequence read quality (i.e., length) and quantity. The sequence obtained by the above two kinds of different sequencing methods were mixed: a) using the software AMOS-3.1.0 minimus2 module, with 454 joining together the results for your reference; b) After assembly and before annotating the transcripts, CD-hit (version 4.6.1) was used to determine whether there was any redundancy in the final data set. The similarity is 95%, and the longest sequence of each category is designated as Unigene.

The RNA-Seq reads from each library were aligned to the cDNAs, and the transcripts were assembled using Bowtie (ver. 0.12.8, single-end mapping method), with the parameters (-v 3 -a --phred64-quals). Expression levels of all unique transcripts mapped onto the full-length cDNAs and contigs were quantified using the RPKM values conducting a correlation analysis of the replicates. The RPKM value of each transcript was calculated using uniquely-aligned reads according to the RPKM formula:

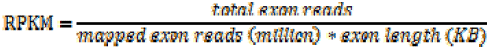

Differentially expressed transcripts were detected using a G-test with an FDR value cut-off < 0.01.

## Transcripts Annotation

Unigene sequences were analyzed by Blastx alignment search (e-value < 1e-5) against protein databases NR, Swiss-Prot, TrEMBL, CDD, PFAM, KEGG and KOG retrieved proteins with the highest sequence similarity (similarity > 30%) for protein functional annotation and classification. In total, almost 90.96% sequences received annotation in this way.

For the gene annotation and functional classification of the reference transcripts set created by pooling raw reads was analyzed by gene ontology (GO) analysis (http://www.geneontology.org/) to understand the distribution of gene functions at the macro level. GO and KEGG functional enrichment analysis were implemented to analyze the transcripts differentially expressed in both wild and mutant wheat types, with differences transcripts being as prospect, while all transcript being as background. Hypergeometric distribution algorithm (phyper) was performed to calculate the prospect transcripts and the P-values of a particular branch of the GO/Pathway classification. Additionally, Fisher’s exact test with false discovery rate (FDR) was used to correct and obtain an adjusted P-value.

### Real-time quantitative reverse transcription PCR

Total RNA of Yunong 201 and Yunong 3114 were extracted from seedlings and seeds, respectively, according to the method of Chen et al. [70] with a slight modification for the extraction buffer containing 0.5 mM Tris-HCl, 0.2 mM NaCl, 1% SDS (w/v), 0.02 mM EDTA, 0.01 mM DTT and 0.07% DEPC-H_2_O (v/v). DNA was removed by digestion with DNAse I (Qiagen, China) before reverse transcription. cDNA first strand was synthesized using M-MLV transcriptase (Invitrogen). Two Steps PrimeScriptTM RT Reagent Kit with gDNA Eraser (Perfect Real Time; TaKaRa) were used for the reverse transcriptase (RT) reactions. The temperature program was adjusted as follows: 2 min at 42 °C, 15 min at 37 °C, 5 s at 85 °C, and then 4 °C. Three biological replicates were performed for each candidate unigene. Specific primers in selected unigenes were designed by software Primer Premier 5.0 and were confirmed by directly sequencing PCR products amplified with the specific primers in Table 1. Amplification with *β*-actin primers [70] was used as an internal control to normalize all data. The relative quantification method (2^ΔΔc^_T_ was used to evaluate quantitative variation between the three replicates.

**Table 1.** Sequencing and draft transcriptome assembly statistics.

|  | All | >=500 | >=1000 | N50 | N90 | Total | Max | Min | Average |
| --- | --- | --- | --- | --- | --- | --- | --- | --- | --- |
|  | (>=200bp) | bp | bp |  |  | Length | Length | Length | Length |
| **Transcript** | 743086 | 414871 | 246268 | 1516 | 382 | 695863802 | 14748 | 201 | 936.45 |
| **Unigene** | 289681 | 89444 | 42109 | 886 | 253 | 172782515 | 14748 | 201 | 596.46 |

The N50 is defined as the length of the contig that spans the median length of the full assembly.

### Alternative splicing analysis

Alternative splicing events and alternatively spliced genes were identified by using BLAST aligning against alternative splicing database, including a common wheat genome sequencing results in NCBI, Chinese spring sequencing results by UK (http://www.cerealsdb.uk.net/cerealgenomics/WheatBP/Documents/DOC_WheatBP.php), Chinese spring sequencing results by French INRA and genome A and D sequencing results reported by Nature [9, 25].

## Results

### Sequencing and assembling of the wheat transcriptomes

Solexa sequencing was performed to obtain transcriptome expressions. Wheat mRNAs from wild type and mutant libraries was sequenced using: a) short-read Illumina solexa technology, and b) long-read 454 GSFLX technology. After quality checks, excluding low quality reads and trimming of adapters and sizes election, 351 million solexa reads (the average length of 90.57 bp) in total were obtained. Trinity was explored for sequence assembly in this experiment.

Each contig included various transcript isoforms, which cause redundancy. To avoid difficulties in a quantitative analysis, the redundant contigs would be removed, by limiting number of contigs and meaning contig length. Thus a non-redundant set of contigs were generated for the next step of assembly program. After removal of redundant, the largest assembly, comprising 14,748 transcripts with an average length of 936.45 bp long and N50 of 1,516 bp (i.e. 50% of the total assembled sequence was present in contigs of this length or longer), was produced by Trinity. The longest transcript of each cluster was definite as Unigene. And 289,681 Unigenes with a mean length of 596.46 bp and N50 of 886 bp were obtained. Finally, we got solexa sequencing assembled transcripts 74,086 sequences, and 454 sequencing assembled transcripts 88,332 sequences (Table 1). We combined the two libraries of sequences for reassembling. The transcripts decreased into 768,141 sequences with average length of 918.74 bp and N50 of 1,498 bp, Unigene of 422,144 sequences with average length of 764.49 bp and N50 of 1267 bp (Table 2).

**Table 2.** Sequencing transcriptome assembly statistics.

|  | All | >=100bp | >=500 | >=1000 | N50 | N90 | Total | Max | Min | Average |
| --- | --- | --- | --- | --- | --- | --- | --- | --- | --- | --- |
|  |  |  | bp | bp |  |  | Length | Length | Length | Length |
| **Transcript** | 768141 | 762662 | 422627 | 247246 | 1498 | 381 | 705722361 | 14748 | 50 | 918.74 |
| **Unigene** | 422144 | 418641 | 188538 | 99839 | 1267 | 306 | 322725389 | 14748 | 50 | 764.49 |

The N50 is defined as the length of the contig that spans the median length of the full assembly.

The transcripts and Unigene expression levels were calculated by the RPKM distribution statistics method (Reads Per kilobase of exon model per Million mapped reads). The sample reads and transcribed sequence were highly correlated, more than 78% of the reads aligned to the transcription assembled sequences. A total of 414,330 Unigenes were expressed, ratio of 98.15%. The distribution of expression values of Unigenes in all the samples existed differences (Max, sd, etc.) (Table 3 and Figure 1).

**Figure 1.**
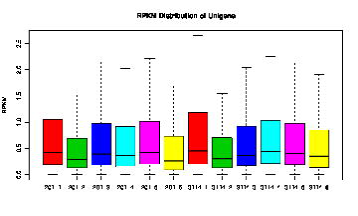
RPKM distribution of Unigenes.

**Table 3.**
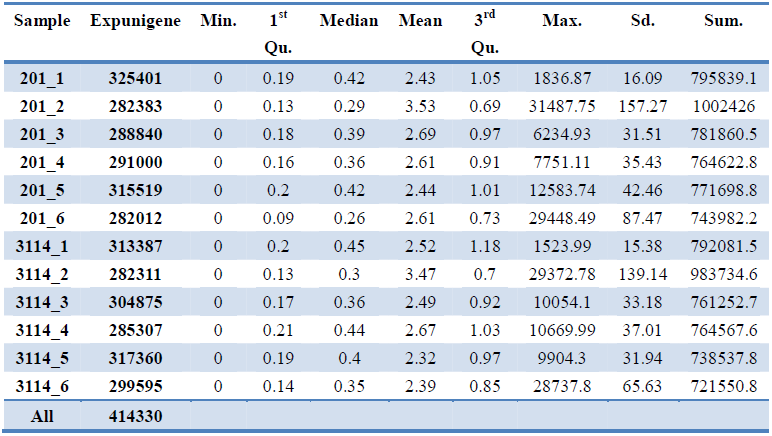
RPKM of Unigenes.

### Functional annotation of differentially expressed genes

To understand the functions of differentially expressed genes, all the Unigenes were aligned by Blastx to the NCBI non-redundant protein database (NR) and the Swiss-Prot, TrEMBL, Cdd, pfam and KOG protein database, using a cut-off E-value of 1e-5.

As for annotation of species distribution, 90.96% of the Unigenes were mainly annotated on the 10 species as showing in Figure 2. 44% of the Unigenes had top matches with sequences from barley (*Hordeum vulgare*), 7% and 3% from rice (*Oryza sativa*), 4% form wheat (*Triticum aestivum*), 4% from sorghum bicolor, 6%, 23% and the rest from others, respectively.

**Figure 2.**
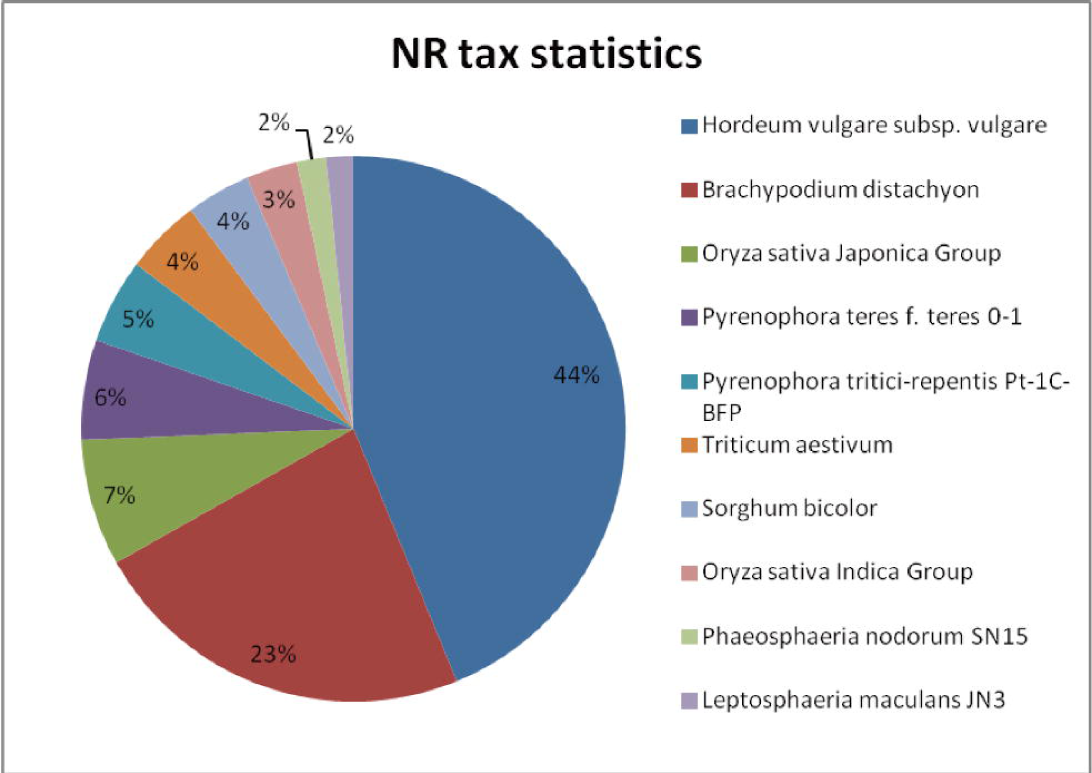
NR annotation of Unigenes. The proportion of Unigenes with matches in NR is exhibited by different colors and annotated beside.

To further evaluate the completeness of this transcriptome library and the effectiveness of the proposed annotation process, we searched the annotated sequences for the genes involved in Clusters of orthologous groups for eukaryotic complete genomes (COG) classifications. Of the 142,575 matches to the nr database, 29,721 sequences have a COG classification (Figure 3). Among the 25 COG categories, the cluster for “signal transduction mechanisms” represents the largest group (6,827, 1.62%), followed by “posttranslational modification, protein turnover, chaperones” (5,290, 1.25%) and “general function prediction only” (4,727, 1.12%). The following categories represent the smallest groups: “cell motility” (5, 0.001%), “Extracellular structures” (80, 0.019%), and “Nuclear structures” (108, 0.026%) (Figure 3).

**Figure 3.**
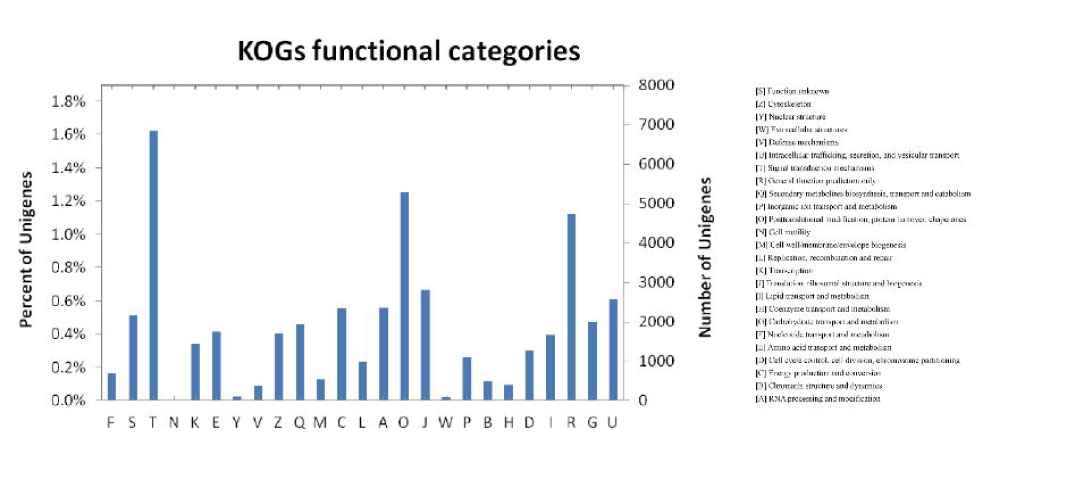
Clusters of orthologous groups (COG) functional classification of Unigenes. 29,721 Unigenes had a COG classification. The y-axis indicates the number of genes in a category. The x-axis indicates the categories of COG functions, represented by capital English characters with annotation on the right side.

Gene ontology (GO) assignments analysis was performed to annotate and classify the gene functions of Unigenes. 89,644 Unigenes were annotated by GO analysis, and all of the annotated Unigenes can be categorized into 64 functional groups (Figure 4). The most abundant categories of GO classification were involved in biological process, cellular components and molecular function, with “cellular process and metabolic processes”, “cell and cell part”, and “binding and catalytic activity” terms gathering the dominant genes in wheat, respectively.

**Figure 4.**
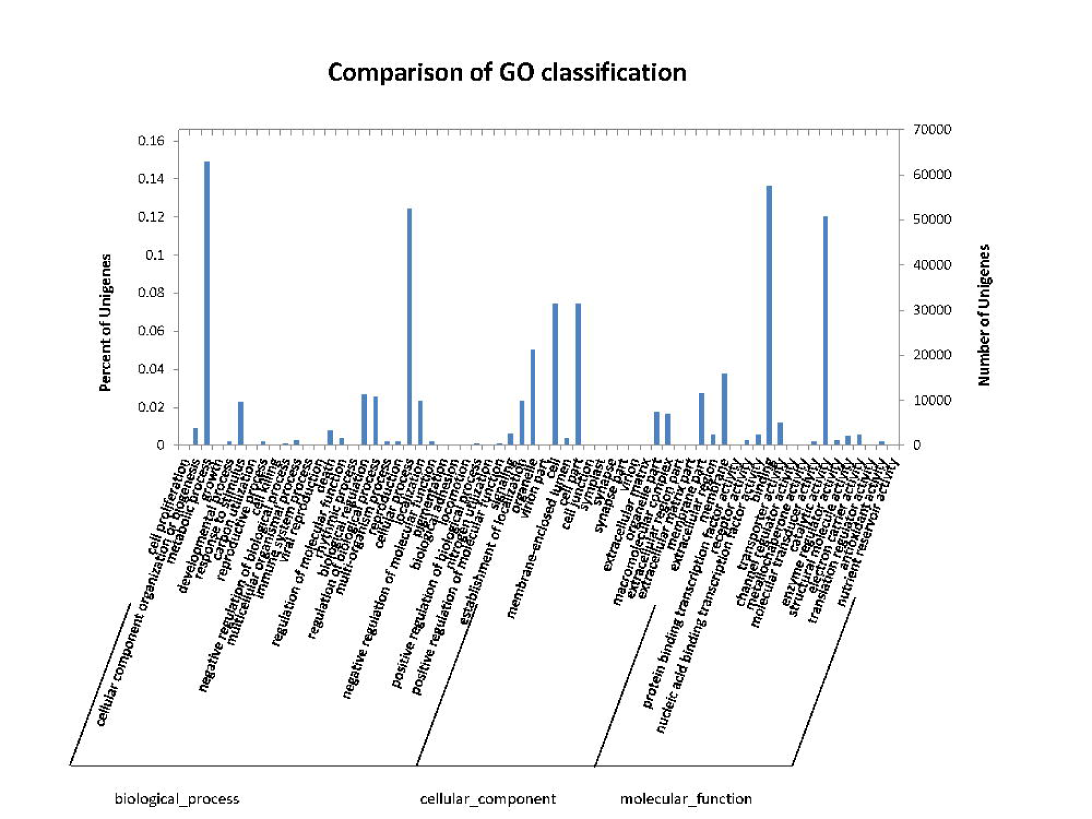
Gene Ontology classification of Unigenes. 89,644 Unigenes were categorized into 64 functional groups of three main categories: biological process, cellular component and molecular function. The y-axis indicates the number of genes in a category.

To characterize the complex biological behaviors for the transcriptome, KEGG (Kyoto Encyclopedia of Genes and Genomes) database was used analyze the pathway annotation of unigenes. In total, 43,501 sequences were assigned to 313 KEGG pathways (Table 4). The pathways with most representation by the unigenes were Metabolism/Carbohydrate Metabolism pathway (1,855), Metabolism/Energy Metabolism pathway (1,767), Genetic Information Processing/Translation pathway (1,309), Genetic Information Processing/Transcription pathway (1,165), Organismal Systems/Environmental Adaptation pathway (1,130).

**Table 4.** The network of Unigenes in KEGG database. 43,501 Unigenes were assigned to 313 KEGG pathways and the top five representative networks were listed.

| Pathway | Pathway_type | Unigene number |
| --- | --- | --- |
| **ko00630, Glyoxylate and dicarboxylate metabolism** | Metabolism/Carbohydrate Metabolism | 1855 |
| **ko00710, Carbon fixation in photosynthetic organisms** | Metabolism/Energy Metabolism | 1767 |
| **ko03010, Ribosome** | Genetic Information Processing/Translation | 1309 |
| **ko03040, Spliceosome** | Genetic Information Processing/Transcription | 1165 |
| **ko04626, Plant-pathogen interaction** | Organismal Systems/Environmental Adaptation | 1130 |

### The difference of gene expression between Yunong 3114 and Yunong 201

Samples clustering showed similar gene expression pattern at different growth stages in Yunong 3114 and Yunong 201 (Figure 5 and Figure S1). As these data showed a large number of genes highly expressed in the initial stage of wheat flowering both in Yunong 3114 and Yunong 201. In contrast, a few clusters showed a little difference (Figure S2).We hypothesized that the treatment of EMS could give birth to some mutant genes or abnormal expression genes, playing a part in the whole lifelong of the plant, which probably resulting in high-yield of Yunong 3114 strain. Therefore, we screened the enrichment GO terms and KEGG pathways for differently expressed genes and pathways in all the six samples of Yunong 3114. The absolute value of “|log2 Ratio|>1” was used as the threshold to identify and compare differentially expressed genes (DEGs) in and between genotypes. Based on the log base 2 fold change values of genes analysis, we condensed and compressed the genes by removing categories that did not show a significantly different change and displaying the categories that did show significant change using a false color heat-map-like display to show up-or down-regulated classes. We identified sixteen enriched specific functional GO categories represented changes in Yunong 3114 compared with Yunong 201: receptor activity, nuclease activity, defense response, recognition of pollen, RNA-directed DNA polymerase activity, endonuclease activity, DNA integration, nucleotidyltransferase activity, transferase activity, protein serine/threonine kinase activity, protein phosphorylation, protein kinase activity, apoptotic process, ATP binding, nucleotide binding and RNA-dependent DNA replication (Table 5). These annotations provided a valuable clue for investigating specific processes, especially those involved in high-yield.

**Figure 5.**
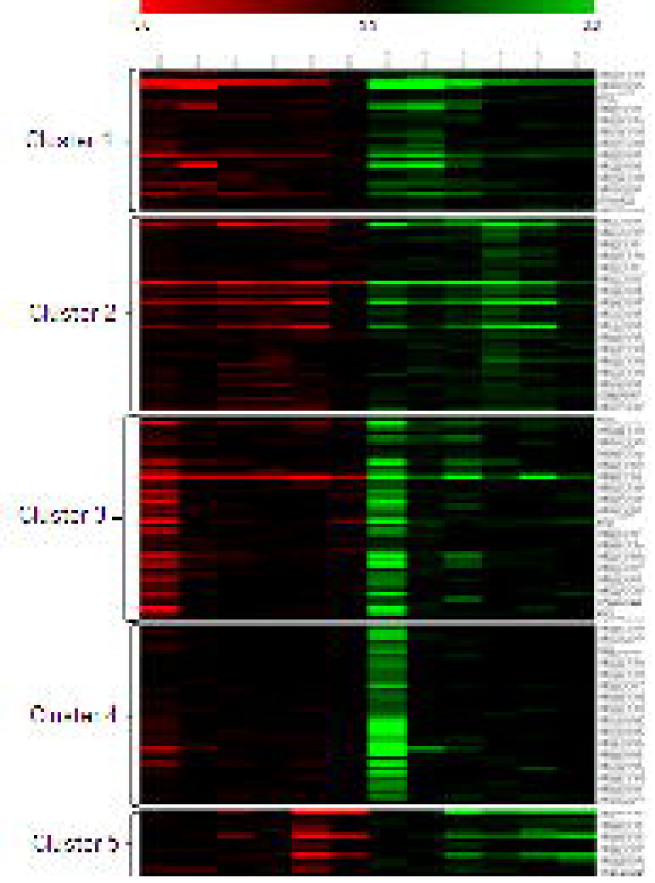
Heat map. The clusters generated based on similarities in expression profile of probe sets differentially expressed in the samples. The red represents samples from wild strain Yunong 201; while the green shows mutant strain Yunong 3114.

**Table 5.**
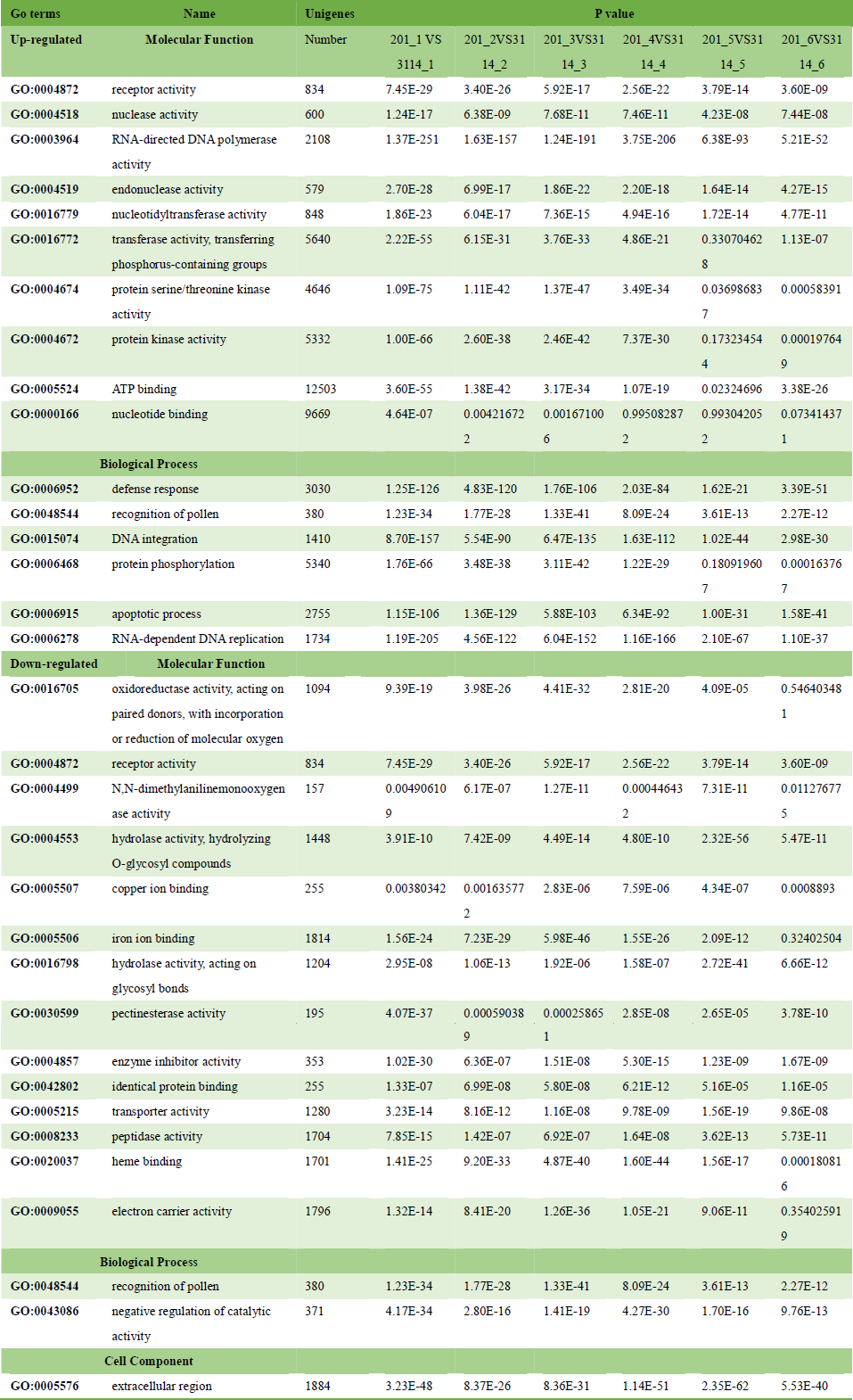
Enriched differently expressed GO terms.

### Recognition of pollen

Pollen, containing the microgametophytes of seed plants, can be recognized by the stigma of an incompatible plant pistils, afterwards pollination and germination to initiate double fertilization. The key pollen factor is a small cysteine-rich protein that interacts directly with the receptor domain of a stigma receptor serine/threonine kinase to initiate haplotype-specific pollen recognition and rejection. We detected 380 unigenes differently expressed in the category of recognition of pollen, of which 94 unigenes significantly up-regulated and 63 down-regulated (Figure 6). Among these 69 of the up-regualted genes and 54 of the down-regulated genes possess annotations in Swissprot database, with encoding proteins 19 and 22 respectively. Most of them encode receptor-like protein kinase (RLK) family proteins, especially a G-type lectin S-receptor-like serine/threonine protein kinase (GsSRK). In addition, in our data, it is interesting to found out that 9 from the sharing 11 proteins were RLK proteins, which were encoded by 41 and 33 unigenes, presented in up-regulated and down-regulated categories respectively (Figure 6).

**Figure 6.**
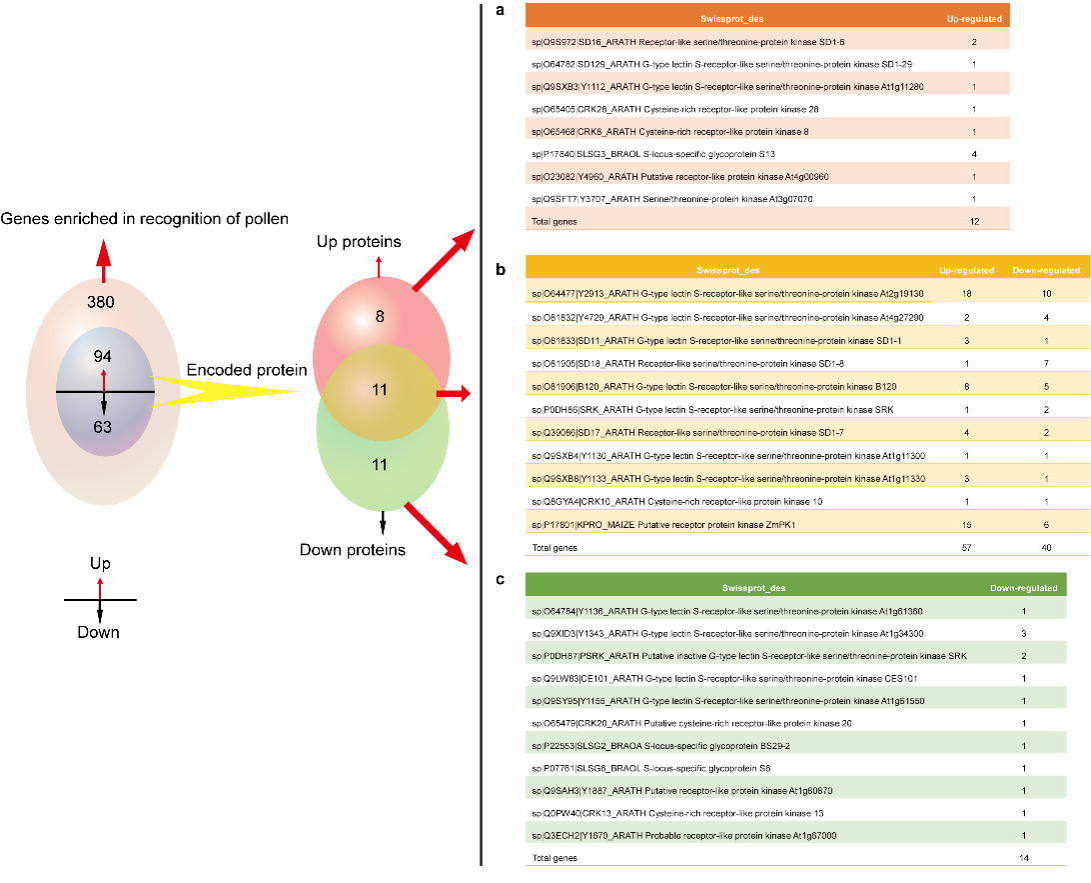
Venn diagram comparison of differentially expressed genes (DEGs) in enrichment GO term of recognition of pollen in Yunong 3114 and 201. The pink cycle in the left represents genes enriched in the category of recognition of pollen; the blue cycle within shows genes of up-regulation and down-regulation. The up or down arrows indicated that DEGs were up-or down-regulated; The red and green ovals show proteins encoded by up-and down-regulated DEGs in the category with annotations in Swissprot, which are listed in the right tables. a) shows proteins only encoded by up-regulated genes; b) proteins encoded by both up-and down-regulated genes; c) proteins encoded by down-regulated genes.

A set of nulli-tetrasomic lines and ditelosomic aneuploids available in the Chinese Spring background were evaluated for physical mapping of 157 up/down-regulated unigenes using PCR and the fragment-specific primers. Results indicated that 10 out of 157 unigenes were located on the chromosome 2AL, 4 of them on 2BL and 10 of them on the 2DL. Relative expression levels of 16 RLK-related unigenes were further confirmed by real-time PCR. Results indicated that 7 and 9 of them were up-regulated and down-regulated in Yunong 3114, which is generally consistent with results analyzed by RNA-Seq.

### Apoptotic process and defense response

Similar with apoptosis of animals, plants undergo programmed cell death (PCD), which plays an important role in the process of development [26], germination of cereal grains and defense against pathogens [27]. PCD is triggered to restrict pathogen growth when a plant disease resistance (R) protein recognizes a corresponding pathogen virulence protein. Our results showed 699 unigenes up-regulated in the enriched GO term of apoptotic process, and 740 unigenes up-regulated in defense response category (Figure 7 upper). As showed in Figure 7, 539 genes in PCD and 465 genes in defense response were non-annotated. While among the genes with annotations, 137 genes are shared between PCD and defense response. In addition, we found few genes in the category of PCD directly encode apoptotic related proteins, instead of a high percentage of genes in the main accumulation profile encoding proteins that belong to disease resistant (R) proteins family (Figure 7 lower). Our results showed that 25 R proteins presented up-regulated. These findings indicate that the differently expressed R genes may play important roles in high-yield phenotype of Yunong 3114.

**Figure 7.**
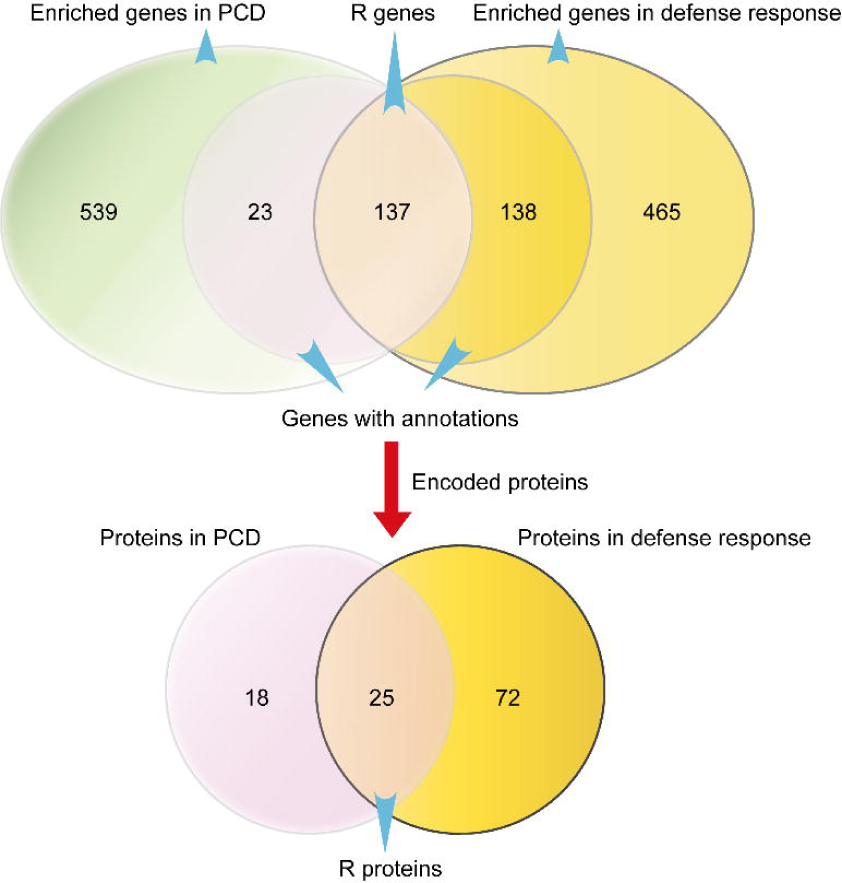
Venn diagram comparison of DEGs in PCD and defense response in Yunong 3114 and 201. The upper cycles represent DEGs enriched in PCD and defense response; the lower cycles show proteins encoded by DEGs in PCD and defense response with annotations in Swissprot. The representation of each cycle is marked by blue arrows.

### Molecular function and other metabolism processes

As mentioned above, of the 89,644 Unigenes, we also found a large amount genes enriched in ATP binding, DNA synthesis, kinase activity and *etc.* (Table 5) which is essential for providing energy for nutrient synthesis and accumulation. It is worth noting that the zeatin biosynthesis pathway, which synthesized a class of phytohormones involved in various processes of growth and development in plants, presented up-regulated in all the stages of strain Yunong 3114, suggesting an underlying relationship with high-yield of Yunong 3114. In addition, we also detected the homologous genes of the known large grain genes in rice. 15 unigenes encoding F-box/Kelch-repeat proteins differently expressed in Yunong 3114, which probably benefit for increasing yield.

### Alternative splicing in wheat strains Yunong 201 and Yunong 3114

Alternative splicing (AS) serves as an important regulatory mechanism in regulating gene expression during development and undergoing different environmental pressure, contributing to the generation of proteomic and functional complexity in higher organisms [24, 28–30]. To identify potential AS events, we carried out computational analyses, mapping the assembled expressed sequence tags to the genome predicted gene regions, to determine all potential AS events. In our results, we identified 11521 and 13331 alternative splicing events based on A genome in wheat Yunong 201 and Yunong 3114 respectively (Table 6). Our study identified that retention of introns is the major type of alternative splicing events (>77%) in both strains. By comparing the AS ratio of wild strain Yunong 201 with mutant strain Yunong 3114, we found that the retained introns (RI, a type of AS) ratio is a little higher in Yunong 3114 with the ratios of other AS types reduced, which indicates the mutation influence AS. Then, we analyzed AS in GsSRK and RLK genes, however, the alternative exon of RLK gene is deferent in wild type and mutant strain, 22276-22358 in Yunong 201, while 22836-22911 in Yunong 3114 (Table 7). We also identified a gene related to recognition of pollen undergoing AS, however, it failed mapping to the Swissprot database. Our results suggest that mutation affected AS, which may play an important role in regulating wheat yield. During the analysis, it is found that AS based on D genome is significant less than AS based on A genome, which may due to the smaller size of wheat D chromosome [31].

**Table 6.**
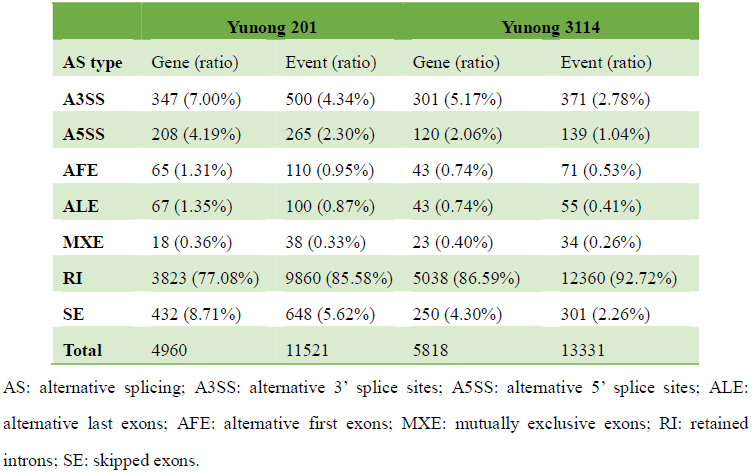
AS events and alternatively spliced genes in wheat Yunong 201 and Yunong 3114.

|  | Yunong 201 |  | Yunong 3114 |  |
| --- | --- | --- | --- | --- |
| AS type | Gene (ratio) | Event (ratio) | Gene (ratio) | Event (ratio) |
| A3SS | 347 (7.00%) | 500 (4.34%) | 301 (5.17%) | 371 (2.78%) |
| A5SS | 208 (4.19%) | 265 (2.30%) | 120 (2.06%) | 139 (1.04%) |
| AFE | 65 (1.31%) | 110 (0.95%) | 43 (0.74%) | 71 (0.53%) |
| ALE | 67 (1.35%) | 100 (0.87%) | 43 (0.74%) | 55 (0.41%) |
| MXE | 18 (0.36%) | 38 (0.33%) | 23 (0.40%) | 34 (0.26%) |
| RI | 3823 (77.08%) | 9860 (85.58%) | 5038 (86.59%) | 12360 (92.72%) |
| SE | 432 (8.71%) | 648 (5.62%) | 250 (4.30%) | 301 (2.26%) |
| Total | 4960 | 11521 | 5818 | 13331 |

AS: alternative splicing; A3SS: alternative 3' splice sites; A5SS: alternative 5' splice sites; ALE: alternative last exons; AFE: alternative first exons; MXE: mutually exclusive exons; RI: retained introns; SE: skipped exons.

**Table 7.** AS events of pollen related genes in wild and mutant strains.

| RefgeneID | Location<br>(strand) | Constitutive<br>exon | Alternative<br>exon | Constitutive<br>exon | Unigene | AS<br>type | Strain | Length | Swissprot_des |
| --- | --- | --- | --- | --- | --- | --- | --- | --- | --- |
| **TRIUR3_25**<br>**324** | scaffold41<br>665 (-) | 23440-23547 | 23440-23550 | 23667-23999 | comp19669<br>9_c0_seq1 | A3SS | Yunong<br>201<br>(wild) | 1273 | sp\|O81906\|B120_ARATH<br>G-type lectin<br>S-receptor-like<br>serine/threonine-protein<br>kinase B120 |
| **TRIUR3_24**<br>**895** | scaffold76<br>660 (-) | 21964-22275 | 22276-22358 | 22359-22509 | comp21116<br>9_c0_seq10 | RI | Yunong<br>201<br>(wild) | 1641 | sp\|Q39086\|SD17_ARATH<br>Receptor-like<br>serine/threonine-protein<br>kinase SD1-7 |
| **TRIUR3_24**<br>**895** | scaffold76<br>660 (-) | 22595-22835 | 22836-22911 | 22912-23122 | comp21116<br>9_c0_seq10 | RI | Yunong<br>3114<br>(mutant) | 1641 | sp\|Q39086\|SD17_ARATH<br>Receptor-like<br>serine/threonine-protein<br>kinase SD1-7 |
| **TRIUR3_02**<br>**437** | scaffold96<br>181 (-) | 38493-39672 | 39673-39749 | 39750-39816 | comp18089<br>7_c0_seq2 | RI | Yunong<br>3114<br>(mutant) | 3045 | NA |

## Discussion

### Transcriptome sequencing and GO annotaion

To the best of our knowledge, transcriptomics is a powerful tool, with larger breadth of the discovery potential over microarrays, to analyze gene expression, single nucleotide polymorphism (SNP) surveys, quantitative trait loci (QTL) studies, genomic scans of diversity, and so forth [32–34]. In order to get a better knowledge of the whole genomic information and the underlying mechanism of high-yield of wheat as well as analysis the relative genes, in the present study we performs the first *de novo* transcriptome sequencing of mixed stages of the new high-yield mutant wheat strain Yunong 3114 by comparing with common strain Yunong 201. The characterization of the wheat transcriptome can help elucidate the molecular mechanisms underlying high-yield of Yunong 3114. Using the Solexa or Roche/454 massively parallel pyrosequencing technology, we have generated 351 million solexa reads. After filtering, these transcripts were *de novo* assembled and extensively annotated. 422,144 Unigenes were obtained, of which 89,644 Unigenes were annotated to 3 terms in the GO database and mapped to 313 KEGG pathways. These results represent the significant expansion of the wheat transcript catalog and provide a comprehensive material basis for future functional and expression analysis of genes of interest.

Gene ontology enrichment analysis is a useful tool to investigate specific processes, unique functions, and pathways in transcriptome research [35]. Based on this consideration, we analyzed the statistically significant enrichment of specific GO terms in the transcripts. In addition, we hypothesized that the mutants caused by EMS functions all the lifelong of Yunong 3114. And the genes accumulated in GO terms with continuing functions or differently expressed may contribute to high-yield of Yunong 3114. Our transcriptome data showed that the differentially expressed genes with consistent expression patterns in both mutant and common wheat strains. However, we found the basal differently expressed genes accumulate in several GO categories, such as regulation of pollen, apoptotic process, defense response, DNA integrate and so forth. Moreover, among the transcripts, GOs associated with ATP binding, nucleotide binding, protein phosphorylation, receptor activity, protein kinase activity and endonuclease activity were significant up-regulated. Plant receptors function in the recognition and transduction of pathogen signals that give rise to reprogramming of cellular metabolism. Endonucleases play a key role in replication, restriction, repair and recombination, as well as in nucleus destruction of PCD in all organisms. It is reported that the functions of various R proteins also required phosphorylation, protein degradation, or specific localization within the host cell [36]. These enrichment GO terms will drive some interesting analysis in the future. Therefore, we focused on several sets of genes in the interesting GO terms compared with the whole transcriptome background to identify genes involved in understanding the gene expression changes in high-yield mutant strain Yunong 3114.

As we known, the key factors involved in high-yield are including grain number, grain size. Several genes’ functions in regulation grain size have been identified in rice. And the complete sequence of the rice genome has provided an extremely useful parallel road map for genetic and genomics studies in wheat. It is demonstrated that the rice genome possess a major grain length quantitative trait locus (QTL), qGL3, which encodes a putative protein phosphatase with Kelch-like repeat domain (OsPPKL1), which is essential for the OsPPKL1 biological function. OsPPKL1 together with its homolog OsPPKL3 are function as negative regulators of grain length, whereas homolog OsPPKL2 as a positive regulator [37]. Another study demonstrated that two novel major genes named GS2 and GS3 involved in the regulation of grain width and length in rice, GW2 for grain width [38], GS3 for grain length [39]. In the present study, we also detected 15 unigenes encoding F-box/Kelch-repeat proteins differently expressed in Yunong 3114. However, whether these genes also play any function in grain length regulation in wheat is worth further investigation.

### RLKs in recognition of pollen and high-yield of wheat

A high percentage of genes in the main accumulation profile also attracted our attentions. One of our most interesting observations was related to genes in the RLK family that encode receptor-like serine/threonine kinase proteins. The cell surface RLK family has been reported to play important roles in various physiological processes, including perceiving external signals [40–42], plant growth and development [43, 44], plant defense responses against pathogens [45, 46], and so on. RLKs are classified into different groups according to the large variability of the extracellular domain organization (over 20 structures exist), such as leucine-rich repeats, S domains, epidermal growth factor-like repeats, and lectin binding domains [47, 48]. Previous studies indicated that G-type RLKs function in self-incompatibility (SI) in flowering plants and plant defense [49–51]. Moreover, a recent research suggested that over expression of GsSRK in Arabidopsis exhibited enhanced salt tolerance and higher yields under salt stress, with higher germination rates, higher chlorophyll content, more green and open leaves, lower ion leakage, higher plant height, and more siliques at the adult developmental stage [52]. This is in accordance with our results that GsSRK is significantly up-regulated in Yunong 3114. We tried to find some clues to reveal the underlying relationship. Therefore, the statistics of locuses of RLK family genes in wheat genome is checked. And we found that 10, 4 and 10 out of 157 unigenes were located on the chromosome 2AL, 2BL and 2DL, respectively. It is suggested that these genes are only one of the reasons for the high yield caused by mutations, rather than the position of the mutation. The discovery of accumulation of receptor-like serine/threonine kinase proteins in Yunong 3114 probably reveals an important involvement with wheat high-yield, which needs further investigation.

In plants, AS functions in a wide range of processes including development, disease resistance and stress response [24, 53, 54], especially in the generation of proteomic and functional diversities [55, 56]. It is reported that serine/arginine-rich proteins, are extensively alternatively spliced [57]. In our results wheat AS-associated genes undergo multiple AS events producing a variety of transcripts from a single gene. These AS events were further classified into seven different types: alternative 3’ splice sites (A3SS), alternative 5’ splice sites (A5SS), alternative last exons (ALE), alternative first exons (AFE), mutually exclusive exons (MXE), retained introns (RI), skipped exons (SE). The different types of AS may reflect underlying differences in pre-mRNA splicing regulation, affect the protein coding sequence or generate unproductive mRNAs to affect transcript levels, which further influence the downstream functions, such as regulating grain weight and length. In our data, RI is the most frequent AS type, this is consistent with previous reports in *Arabidopsis* and Rice [57–59]. However, we found that RI type of AS event ratio is more prevalent in mutant strain, which may result in different levels of transcripts with wild strain. Moreover, our results showed that RLK genes significantly up-regulated in mutant strain, we deducted that these differentially expressed genes in the two genotypes were due to the fragment deletion on the distal part of transcripts or AS in the mutant strain, which may increase protein variation and complexity. Interestingly, our AS analysis showed AS event in GsSRK and RLK genes, firmly supporting our speculation. Furthermore, the AS of these genes probably helps with regulating transcripts levels illustrating the extremely high complexity of transcriptome regulation affecting development.

### R genes in PCD and defense response may be conducive to high-yield of wheat

Another set of genes we are interested in is involved in PCD and defense response. PCD process, which programmed eliminated unwanted cells, is an essential process of development of animals and plants. While in plant cells, it possesses unique characteristics, such as presence of chloroplasts, a prominent vacuole, and the cell wall, which affect PCD [60]. It is an essential process occurring throughout the plant’s life cycle, such as grain germination in initial stages of grain development [61, 62], fertilization, xylogenesis, and senescence [63, 64], and also in the accumulation of storage compounds of starchy endosperm during maturation [65, 66]. Mounting researches suggested that PCD and disease resistance are complicated linked in plants, in virtue of the simultaneous activation of disease resistance and cell death upon pathogen recognition by disease resistance (R) proteins [67]. Plant disease resistance is often activated by genes with nucleotide binding site (NBS) and leucine-rich repeat (LRR) or serine/threonine protein kinase (S/TPK) domains [68]. The transcriptome data show patterns of expression of different genes involved in PCD are similar. We found numerous genes enriched in PCD category encode disease resistance proteins. Previous studies have identified that a direct effect of an R gene on yield implies an underlying mechanistic relationship. A few genes involved in or related to diseases resistant incurred a yield penalty. However, some others are not. For instance, the 1RS chromosome arm from rye, carrying several disease resistance genes, is associated with increased production even in the absence of disease [69]. In our data, we identified several enriched expressed R genes up-regulated in Yunong 3114 against Yunong 201, which are possibly critical for in Yunong 3114. As indicated previously, the up-regulated R genes in Yunong 3114 would reinforce the ability of disease resistance, more importantly, they may increase yield even during no disease conditions. Unfortunately, we found that the R genes were distributed in each chromosome with no obvious distribution characteristics. Furthermore, many reports suggest that alternative transcripts may limit the expression of R proteins or encode truncated R proteins with a positive role in defense activation. In addition, R gene alternative splicing is dynamic during the defense response. Thus in our study, we also analyzed AS of R genes as well as RLKs. Interesting, different from RLKs, we detected no AS in R genes. This is probably due to incomplete annotations. However, the further investigation will be addressed in the further studies.

In summary, in this work, we performed a transcriptome sequencing of a new mutant wheat strain Yunong 3114. The functional annotation of the transcriptomes provide a further insight into the gene content, biological processes, molecular functions and pathways conserved in wheat. By comparing with its common wheat strain Yunong 201, we found several enriched GO terms and genes probably associated with high-yield phenotype of Yunong 3114, which are worthy discovered to improve yield for increased food or feed productivity. Our data and results provide essential information for future genetic improvements and management in increase-yield and breeding at the transcriptome level.

## Acknowledgments

This project was funded by the National Natural Science Foundation (31370031), 973 project (2014CB138105) and Program for New Century Excellent Talents in University (NCET-13-0776) of China.

F.C. and D.C. designed the project and performed RNA-seq experiments. Z.D., N.Z. and X.Z. performed the computational analyses. F.C. wrote the paper.

## References

1. Cordain, L., Cereal grains: humanity’s double-edged sword. World Rev Nutr Diet, 1999. 84: p. 19–73.

2. *Food and Agriculture Organization of the United Nations*. [http://faostat.fao.org], 2011. *figures*.

3. Tester, M. and P. Langridge, Breeding technologies to increase crop production in a changing world. Science, 2010. 327(5967): p. 818–22.

4. Schreiber, A.W., et al., Transcriptome-scale homoeolog-specific transcript assemblies of bread wheat. BMC Genomics, 2012. 13: p. 492.

5. Tasleem-Tahir, A., et al., Proteomic analysis of peripheral layers during wheat (Triticum aestivum L.) grain development. Proteomics, 2011. 11(3): p. 371–9.

6. Gao, F., M.C. Jordan, and B.T. Ayele, Transcriptional programs regulating seed dormancy and its release by after-ripening in common wheat (Triticum aestivum L.). Plant Biotechnol J, 2012. 10(4): p. 465–76.

7. Liu, Y., et al., The Relationship between Polyamines and Hormones in the Regulation of Wheat Grain Filling. PLoS One, 2013. 8(10): p. e78196.

8. Berkman, P.J., et al., Next-generation sequencing applications for wheat crop improvement. Am J Bot, 2012. 99(2): p. 365–71.

9. Brenchley, R., et al., Analysis of the bread wheat genome using whole-genome shotgun sequencing. Nature, 2012. 491(7426): p. 705–10.

10. Wan, Y., et al., Transcriptome analysis of grain development in hexaploid wheat. BMC Genomics, 2008. 9: p. 121.

11. Xiao, J., et al., Transcriptome-based discovery of pathways and genes related to resistance against Fusarium head blight in wheat landrace Wangshuibai. BMC Genomics, 2013. 14: p. 197.

12. Krasileva, K.V., et al., Separating homeologs by phasing in the tetraploid wheat transcriptome. Genome Biol, 2013. 14(6): p. R66.

13. Garnica, D.P., et al., Strategies for Wheat Stripe Rust Pathogenicity Identified by Transcriptome Sequencing. PLoS One, 2013. 8(6): p. e67150.

14. Xin, M., et al., Transcriptome comparison of susceptible and resistant wheat in response to powdery mildew infection. Genomics Proteomics Bioinformatics, 2012. 10(2): p. 94–106.

15. Tenea, G.N., F. Cordeiro Raposo, and A. Maquet, Comparative transcriptome profiling in winter wheat grown under different agricultural practices. J Agric Food Chem, 2012. 60(44): p. 10970–8.

16. Winfield, M.O., et al., Plant responses to cold: Transcriptome analysis of wheat. Plant Biotechnol J, 2010. 8(7): p. 749–71.

17. Spannagl, M., et al., Analysing complex Triticeae genomes - concepts and strategies. Plant Methods, 2013. 9(1): p. 35.

18. Kumar, A., et al., Physical mapping resources for large plant genomes: radiation hybrids for wheat D-genome progenitor Aegilops tauschii. BMC Genomics, 2012. 13: p. 597.

19. Li, W., et al., Sequence composition, organization, and evolution of the core Triticeae genome. Plant J, 2004. 40(4): p. 500–11.

20. Tanaka, T., et al., Next-Generation Survey Sequencing and the Molecular Organization of Wheat Chromosome 6B. DNA Res, 2013.

21. Garg, R., et al., Gene discovery and tissue-specific transcriptome analysis in chickpea with massively parallel pyrosequencing and web resource development. Plant Physiol, 2011. 156(4): p. 1661–78.

22. Duan, J., et al., Optimizing de novo common wheat transcriptome assembly using short-read RNA-Seq data. BMC Genomics, 2012. 13: p. 392.

23. Oono, Y., et al., Characterisation of the wheat (Triticum aestivum L.) transcriptome by de novo assembly for the discovery of phosphate starvation-responsive genes: gene expression in Pi-stressed wheat. BMC Genomics, 2013. 14: p. 77.

24. Blencowe, B.J., Alternative splicing: new insights from global analyses. Cell, 2006. 126(1): p. 37–47.

25. Ling, H.Q., et al., Draft genome of the wheat A-genome progenitor Triticum urartu. Nature, 2013. 496(7443): p. 87–90.

26. Kuriyama, H. and H. Fukuda, Developmental programmed cell death in plants. Curr Opin Plant Biol, 2002. 5(6): p. 568–73.

27. Watanabe, N. and E. Lam, Recent advance in the study of caspase-like proteases and Bax inhibitor-1 in plants: their possible roles as regulator of programmed cell death. Mol Plant Pathol, 2004. 5(1): p. 65–70.

28. Black, D.L., Mechanisms of alternative pre-messenger RNA splicing. Annu Rev Biochem, 2003. 72: p. 291–336.

29. Matlin, A.J., F. Clark, and C.W. Smith, Understanding alternative splicing: towards a cellular code. Nat Rev Mol Cell Biol, 2005. 6(5): p. 386–98.

30. Reddy, A.S., Alternative splicing of pre-messenger RNAs in plants in the genomic era. Annu Rev Plant Biol, 2007. 58: p. 267–94.

31. Saintenac, C., et al., Sequence-based mapping of the polyploid wheat genome. G3 (Bethesda), 2013. 3(7): p. 1105–14.

32. Nagaraj, S.H., et al., ESTExplorer: an expressed sequence tag (EST) assembly and annotation platform. Nucleic Acids Res, 2007. 35(Web Server issue): p. W143–7.

33. Bouck, A., S.R. Wessler, and M.L. Arnold, QTL analysis of floral traits in Louisiana iris hybrids. Evolution, 2007. 61(10): p. 2308–19.

34. Kota, R., et al., Snipping polymorphisms from large EST collections in barley (Hordeum vulgare L.). Mol Genet Genomics, 2003. 270(1): p. 24–33.

35. Garg, R., et al., Deep Transcriptome Sequencing of Wild Halophyte Rice, Porteresia coarctata, Provides Novel Insights into the Salinity and Submergence Tolerance Factors. DNA Res, 2013.

36. Martin, G.B., A.J. Bogdanove, and G. Sessa, Understanding the functions of plant disease resistance proteins. Annu Rev Plant Biol, 2003. 54: p. 23–61.

37. Zhang, X., et al., Rare allele of OsPPKL1 associated with grain length causes extra-large grain and a significant yield increase in rice. Proc Natl Acad Sci U S A, 2012. 109(52): p. 21534–9.

38. Song, X.J., et al., A QTL for rice grain width and weight encodes a previously unknown RING-type E3 ubiquitin ligase. Nat Genet, 2007. 39(5): p. 623–30.

39. Fan, C., et al., GS3, a major QTL for grain length and weight and minor QT L for grain width and thickness in rice, encodes a putative transmembrane protein. Theor Appl Genet, 2006. 112(6): p. 1164–71.

40. Stone, J.M. and J.C. Walker, Plant protein kinase families and signal transduction. Plant Physiol, 1995. 108(2): p. 451–7.

41. Lease, K., E. Ingham, and J.C. Walker, Challenges in understanding RLK function. Curr Opin Plant Biol, 1998. 1(5): p. 388–92.

42. Zan, Y., et al., Genome-wide identification, characterization and expression analysis of populus leucine-rich repeat receptor-like protein kinase genes. BMC Genomics, 2013. 14: p. 318.

43. Hematy, K. and H. Hofte, Novel receptor kinases involved in growth regulation. Curr Opin Plant Biol, 2008. 11(3): p. 321–8.

44. Nibau, C. and A.Y. Cheung, New insights into the functional roles of CrRLKs in the control of plant cell growth and development. Plant Signal Behav, 2011. 6(5): p. 655–9.

45. Ederli, L., et al., The Arabidopsis thaliana cysteine-rich receptor-like kinase CRK20 modulates host responses to Pseudomonas syringae pv. tomato DC3000 infection. J Plant Physiol, 2011. 168(15): p. 1784–94.

46. Yang, Y., et al., Proteomic study participating the enhancement of growth and salt tolerance of bottle gourd rootstock-grafted watermelon seedlings. Plant Physiol Biochem, 2012. 58: p. 54–65.

47. Zhao, J., et al., A receptor-like kinase gene (GbRLK) from Gossypium barbadense enhances salinity and drought-stress tolerance in Arabidopsis. BMC Plant Biol, 2013. 13: p. 110.

48. Shiu, S.H. and A.B. Bleecker, Receptor-like kinases from Arabidopsis form a monophyletic gene family related to animal receptor kinases. Proc Natl Acad Sci U S A, 2001. 98(19): p. 10763–8.

49. Sherman-Broyles, S., et al., S locus genes and the evolution of self-fertility in Arabidopsis thaliana. Plant Cell, 2007. 19(1): p. 94–106.

50. Kim, H.S., et al., An S-locus receptor-like kinase plays a role as a negative regulator in plant defense responses. Biochem Biophys Res Commun, 2009. 381(3): p. 424–8.

51. Kim, S., et al., An atypical soybean leucine-rich repeat receptor-like kinase, GmLRK1, may be involved in the regulation of cell elongation. Planta, 2009. 229(4): p. 811–21.

52. Sun, X.L., et al., GsSRK, a G-type lectin S-receptor-like serine/threonine protein kinase, is a positive regulator of plant tolerance to salt stress. J Plant Physiol, 2013. 170(5): p. 505–15.

53. Macknight, R., et al., Functional significance of the alternative transcript processing of the Arabidopsis floral promoter FCA. Plant Cell, 2002. 14(4): p. 877–88.

54. Gassmann, W., Alternative splicing in plant defense. Curr Top Microbiol Immunol, 2008. 326: p. 219–33.

55. Zhang, X.C. and W. Gassmann, Alternative splicing and mRNA levels of the disease resistance gene RPS4 are induced during defense responses. Plant Physiol, 2007. 145(4): p. 1577–87.

56. Guo, S., et al., Transcriptome sequencing and comparative analysis of cucumber flowers with different sex types. BMC Genomics, 2010. 11: p. 384.

57. Palusa, S.G., G.S. Ali, and A.S. Reddy, Alternative splicing of pre-mRNAs of Arabidopsis serine/arginine-rich proteins: regulation by hormones and stresses. Plant J, 2007. 49(6): p. 1091–107.

58. Zhang, G., et al., Deep RNA sequencing at single base-pair resolution reveals high complexity of the rice transcriptome. Genome Res, 2010. 20(5): p. 646–54.

59. E, Z., L. Wang, and J. Zhou, Splicing and alternative splicing in rice and humans. BMB Rep, 2013. 46(9): p. 439–47.

60. Williams, B. and M. Dickman, Plant programmed cell death: can’t live with it; can’t live without it. Mol Plant Pathol, 2008. 9(4): p. 531–44.

61. Dominguez, F. and F.J. Cejudo, Identification of a nuclear-localized nuclease from wheat cells undergoing programmed cell death that is able to trigger DNA fragmentation and apoptotic morphology on nuclei from human cells. Biochem J, 2006. 397(3): p. 529–36.

62. Dominguez, F., J. Moreno, and F.J. Cejudo, The nucellus degenerates by a process of programmed cell death during the early stages of wheat grain development. Planta, 2001. 213(3): p. 352–60.

63. Greenberg, J.T., Programmed cell death: a way of life for plants. Proc Natl Acad Sci U S A, 1996. 93(22): p. 12094–7.

64. Bai, S., et al., Proteomic analysis of pollination-induced corolla senescence in petunia. J Exp Bot, 2010. 61(4): p. 1089–109.

65. Young, T.E., D.R. Gallie, and D.A. DeMason, Ethylene-Mediated Programmed Cell Death during Maize Endosperm Development of Wild-Type and shrunken2 Genotypes. Plant Physiol, 1997. 115(2): p. 737–751.

66. Young, T.E. and D.R. Gallie, Programmed cell death during endosperm development. Plant Mol Biol, 2000. 44(3): p. 283–301.

67. Yang, H., et al., The Arabidopsis BAP1 and BAP2 genes are general inhibitors of programmed cell death. Plant Physiol, 2007. 145(1): p. 135–46.

68. Faris, J.D., et al., A unique wheat disease resistance-like gene governs effector-triggered susceptibility to necrotrophic pathogens. Proc Natl Acad Sci U S A, 2010. 107(30): p. 13544–9.

69. Brown, J.K., Yield penalties of disease resistance in crops. Curr Opin Plant Biol, 2002. 5(4): p. 339–44.

70. Chen, F., Gao, M.X., Zhang, J.H., Zuo, A.H., Shang, X.L., Cui, D.Q. Molecular characterization of vernalization and response genes in bread wheat from the Yellow and Huai valley of China. BMC Plant Biology 2013, 13:199

